# Systematic transcriptomics analysis of calorie restriction and rapamycin unveils their synergistic interaction in prolonging cellular lifespan

**DOI:** 10.1101/2023.10.26.564115

**Authors:** Yizhong Zhang, Arshia Naaz, Trishia Yi Ning Cheng, Jovian Jing Lin, Mohammad Alfatah

**Author notes:** To whom the correspondence should be addressed. (Mohammad Alfatah).

## Abstract

Aging is a multifaceted biological process marked by the decline in both mitotic and postmitotic cellular function, often central to the development of age-related diseases. In the pursuit of slowing or even reversing the aging process, a prominent strategy of significant interest is calorie restriction (CR), also known as dietary restriction, and the potential influence of a drug called rapamycin (RM). Both CR and RM have demonstrated the capacity to extend healthspan and lifespan across a diverse array of species, including yeast, worms, flies, and mice. Nevertheless, their individual and combined effects on mitotic and postmitotic cells, as well as their comparative analysis, remain areas that demand a thorough investigation. In this study, we employ RNA-sequencing methodologies to comprehensively analyze the impact of CR, RM, and their combination (CR+RM) on gene expression in yeast cells. Our analysis uncovers distinctive, overlapping, and even contrasting patterns of gene regulation, illuminating the unique and shared effects of CR and RM. Most notably, our findings reveal a synergistic effect of CR+RM in extending the lifespan of postmitotic cells, a result validated in both yeast and human cells. This research offers valuable insights into the processes of aging and presents potential strategies for enhancing healthspan and delaying the onset of age-related diseases. These findings have the potential to revolutionize our approach to implementing these interventions under specific conditions and within the context of age-related diseases.

## INTRODUCTION

Age-related diseases, often linked to dysfunctional mitotic and postmitotic cells, encompass a range of conditions ^1–3^. These include cancer, where unchecked mitotic cell division leads to the formation of tumors ^4^; neurodegenerative diseases, such as Alzheimer’s, Parkinson’s, and Huntington’s disease, characterized by dysfunction and death of postmitotic neurons ^5^; cardiovascular ailments, where dysfunction in both mitotic and postmitotic cells contributes to conditions like atherosclerosis, arrhythmias, and heart failure ^6^; muscle atrophy, seen in sarcopenia, resulting from the deterioration of postmitotic muscle cells; osteoporosis, driven by age-related changes in postmitotic bone cells, leading to decreased bone density and an increased risk of fractures ^7^; diabetes, particularly Type 2 diabetes, stemming from the dysfunction of postmitotic pancreatic beta cells ^8^; age-related macular degeneration, affecting the macula in the retina and leading to vision loss, primarily involving postmitotic cells in the retina ^9^; and skin diseases, including age-related skin conditions that manifest as wrinkles, sagging, and other dermatological issues, primarily involving postmitotic skin cells ^10^.

Aging research has assumed a prominent role as the aging population experiences rapid growth, underscoring the urgent imperative to confront aging-associated diseases. The pursuit of geroprotection strategies, which encompasses interventions like exercise, dietary manipulation, and anti-aging supplements, is gaining momentum in this context. Notably, calorie restriction and the use of rapamycin have emerged as promising approaches, demonstrating their efficacy in extending lifespan across various species ^11^. These strategies offer potential avenues to mitigate the impact of aging-associated diseases, ultimately extending healthspan and enhancing the quality of life for individuals as they age.

Calorie restriction (CR), a dietary regimen characterized by reduced caloric intake, has exhibited the ability to enhance both maximum and median lifespans across a wide spectrum of organisms while delaying the onset of age-related pathologies. This effect appears to be evolutionarily conserved, spanning diverse taxa such as single-celled organisms, invertebrates, and vertebrates ^12–16^. These mechanisms largely converge on nutrient signaling pathways, including AMP-activated protein kinase (AMPK) and the Target of Rapamycin Complex 1 (TORC1) signaling. Consequently, these pathways modulate cellular processes such as autophagy, mitochondrial function, oxidative stress responses, and protein synthesis, collectively influencing healthspan and survival outcomes. The positive effects of dietary interventions have remained evident even in instances of macronutrient alteration, specific dietary component restrictions (e.g., methionine), and variations in feeding schedules including periodic fasting. Remarkably, recent human trials involving CR demonstrated improved thymic function and anti-inflammatory effects on adipose tissues through a 14% reduction in calorie intake over a span of two years ^w17^. However, conflicting findings have emerged, with the effectiveness of CR differing among experiments and species. For instance, CR has failed to extend the lifespan in specific species, such as flies, nematodes, certain rodent strains, and rhesus monkeys ^18^. Notably, initiation of CR at an advanced age has even been associated with increased mortality in rodent and primate models^19,20^.

Rapamycin initially discovered as an antifungal agent in 1960, has since evolved into a multi-functional compound ^21^. Its applications range from immunosuppression for organ transplant recipients to cancer chemotherapy targeting cell proliferation ^22,23^. Of particular interest is rapamycin’s role as a TORC1 inhibitor, which simulates a state akin to calorie restriction, effectively promoting health improvements across diverse animal models and inhibiting cellular senescence. This process involves rapamycin forming a complex with the yeast FK506 binding protein, FKBP, resulting in the destabilization and subsequent inhibition of TORC1 ^24^. The TORC1 signaling pathway’s conservation spans a broad array of species. Genetic and pharmacological inhibition of TORC1 activity has shown the potential to extend both lifespan and healthspan across budding yeast, nematodes, flies, mice, canines, and primates ^22,23,32–36,24–31^. Despite rapamycin’s demonstrated potential in extending lifespan and mitigating age-related diseases, certain age-associated traits remain unaffected or even exacerbated ^37^. Moreover, there is evidence suggesting rapamycin’s involvement in negative outcomes, including hyperglycaemia, glucose intolerance, insulin resistance, and hyperlipidaemia ^38–41^. The mechanisms underpinning rapamycin’s dual effects remain elusive.

In light of the complexities of aging biology, most discoveries in this field have been made through studies on model organisms with short lifespans, including *Saccharomyces cerevisiae*, a budding yeast. *S. cerevisiae* exhibits conserved pathways relevant to human aging and age-related diseases, encompassing nutrient signaling and mitochondrial homeostasis ^42,43^. The aging phenomena observed in yeast, both replicative and chronological, bear a striking resemblance to human mitotic and postmitotic cellular aging, respectively ^44^. Yeast exponential dividing cells mimic human mitotic cellular aging, while yeast stationary phase non-dividing cells correspond to human postmitotic cellular aging.

Yeast mitotic cells divide exponentially under nutrient-replete conditions, exclusively utilizing glucose as the main carbon source for growth. However, upon nutrient deprivation in the culture medium, yeast enters the stationary phase, ceasing division, and non-dividing postmitotic cells remain viable ^11,45,46^. This transition into the stationary phase is a progressive cellular process coupled with metabolic adaptation. When glucose is exhausted in the culture medium, yeast cells undergo a transition from fermentative to respiratory metabolism, known as the diauxic shift, followed by the onset of the stationary phase ^47,48^. The depletion of both glucose and ethanol is associated with cell cycle arrest, marking the transition between mitotic and postmitotic cellular conditions. Consequently, the responses to anti-aging interventions can substantially differ between the mitotic and postmitotic growth conditions. Utilizing these dynamic changes in yeast model organisms could potentially provide an answer to the diverse treatment results observed with calorie restriction and rapamycin, which may be dependent on metabolic and cell type factors.

This study harnesses the RNA-sequencing approaches and embarks on a comprehensive timelapse experiment, precisely observing both mitotic and postmitotic yeast cells. The primary objective is to delve into the intricate transcriptomic responses that arise when yeast cells are subjected to distinct interventions: calorie restriction (CR), rapamycin (RM), and the intriguing combination of both (CR+RM). This multifaceted approach enables us to provide a realistic picture of how these interventions individually and in combination influence the processes underlying cellular aging. By scrutinizing the transcriptomic changes across these different conditions, we seek to uncover the complex molecular mechanisms that drive cellular aging. Are there unique pathways activated by CR that differ from those triggered by RM? How does the combination of these interventions impact gene expression patterns compare to their individual effects? By dissecting the intricate interplay between CR, RM, and their combined effects on yeast cells, we hope to shed entirely new and illuminating light on the potential anti-aging benefits of these interventions. Such insights hold the promise of not only advancing our understanding of the fundamental biology of aging but also paving the way for innovative strategies in treatments and drug development that may extend the healthy lifespan of individuals.

## RESULTS

### Transcriptomic signatures in mitotic and postmitotic cellular growth conditions

In yeast model studies, amino acid auxotrophic *S. cerevisiae* strains are frequently employed, notwithstanding the pronounced impact of amino acid nutrients on various biological processes, such as cellular aging and transcriptomics ^49,50^. In our investigation, we utilized prototrophic strain CEN.PK113-7D ^51^ to exclude any potential extraneous bias originating from amino acid constituents or levels in the culture medium. To evaluate the transcriptomic profiles of both mitotic cells (MiC) and postmitotic cells (PoMiC) of yeast, we conducted a series of experiments under controlled laboratory conditions. Yeast cells were cultured in a synthetic defined (SD) medium comprising yeast nitrogen base with ammonium sulfate, devoid of amino acids, and supplemented with 2% glucose as the carbon source. The sampling occurred at two distinct time points: 5 hours into the mitotic growth (exponential phase) and 72 hours (stationary phase), corresponding to the postmitotic growth (Figures. 1a and 1b). These time points were selected following inoculation of overnight-grown yeast cultures, which were diluted to an optical density (OD600nm) of 0.2, into fresh SD medium.

**Figure 1.**
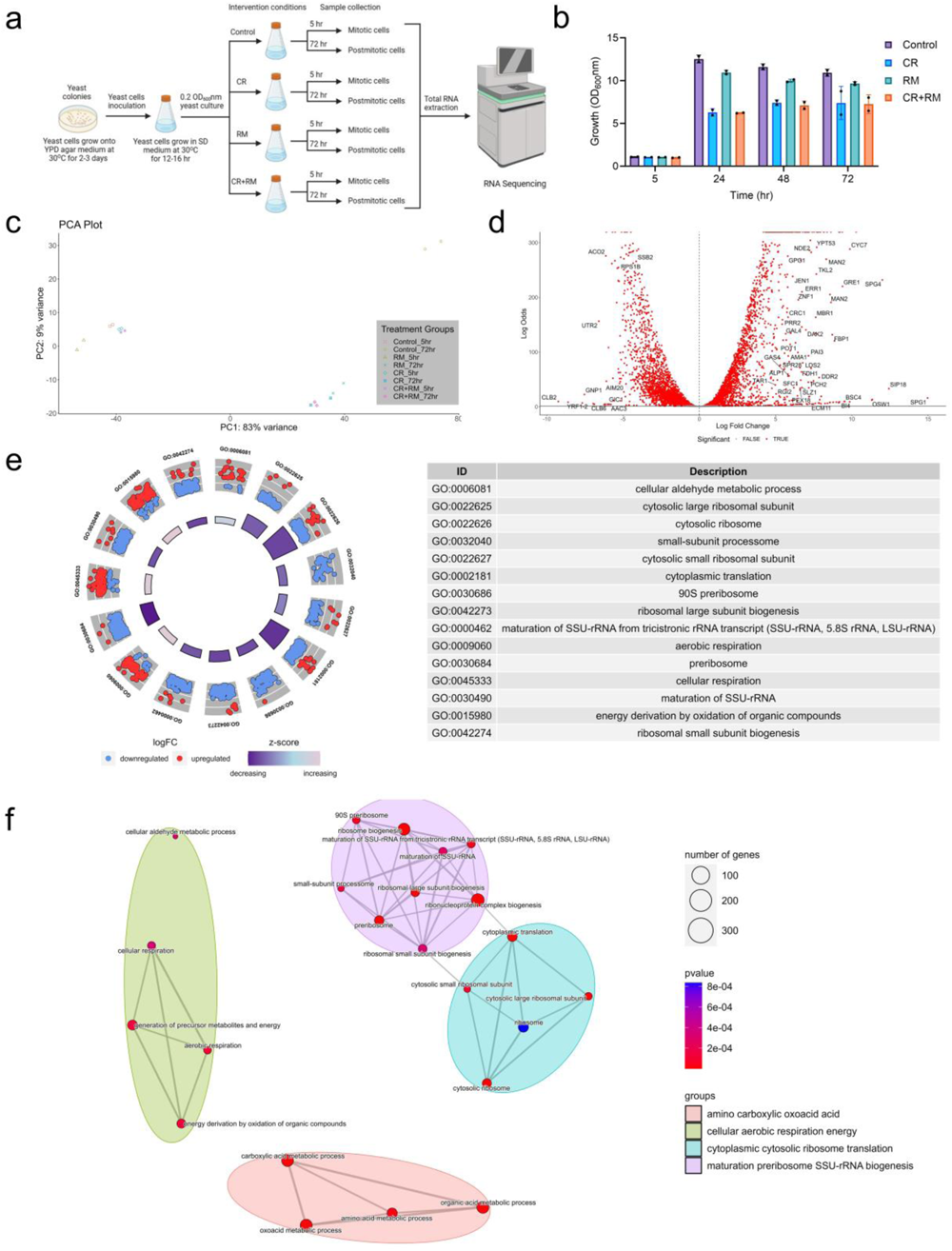
Transcriptomic analysis of mitotic and postmitotic cells under intervention conditions. (a) Schematic of the study workflow. The control treatment used SD medium comprising yeast nitrogen base with ammonium sulfate without amino acids and supplemented with 2% glucose. Calorie restriction (CR) employed SD medium with 0.5% glucose, rapamycin (RM) utilized SD medium with 2% glucose and 2 nM rapamycin, and combined calorie restriction and rapamycin (CR+RM) interventions applied SD medium with 0.5% glucose and 2 nM rapamycin. An equal volume of DMSO solvent is added to normalize the samples to the rapamycin-treated samples. (b) Growth assay of the prototrophic wild-type *Saccharomyces cerevisiae* CEN.PK113-7D strain under control, CR, RM, and CR+RM treatments. Growth optical density (OD_600_nm) was measured at 5, 24, 48, and 72 hours after the initial inoculation with an overnight-grown culture. (c) Principal Component Analysis (PCA) plot showcasing the distribution and clustering of all samples analysed in this study. (d) Volcano plots depicting the distribution of differentially expressed genes (DEGs) in control sample in stationary phase postmitotic cells compared to exponential phase mitotic cells. (e) Circular plots presenting the top gene ontology (GO) enrichments associated with DEGs. (f) Network diagrams visually depict the enriched gene ontologies organized into distinct functional groups.

Three distinct treatment groups were established for comparative analysis; (i) Calorie restriction (CR) group: Yeast cells were cultivated under conditions featuring 0.5% glucose, mimicking a calorie-restricted environment, (ii) Rapamycin (RM) group: Yeast cultures were inoculated with 2 nM rapamycin, a specific inhibitor of the TORC1 pathway and (iii) Combined treatment (CR+RM) group: Yeast cells were exposed to both calorie restriction (CR) and rapamycin (RM) treatments concurrently. A vehicle control group was also included, wherein cells were treated with the DMSO solvent. Rigorous efforts were undertaken to ensure the uniformity of various experimental parameters, including treatment duration, treatment concentrations, and environmental conditions such as incubation temperature and the aeration of culture flasks, across all experimental groups. These measures were implemented to minimize potential confounding factors and ensure the robustness of our transcriptomic analyses.

Principal Component Analysis (PCA) reveals that the transcriptomic profiles of control samples during the exponential and stationary phases distinctly separate along PC1 and PC2 (Figure. 1c). This result highlights significant differences in gene expression profiles between MIC and PoMiC growth conditions. During the stationary phase, 1982 genes were upregulated (fold change ≥ 1), while 1960 genes were downregulated (fold change ≤ -1) in comparison to the exponential phase (Figure. 1d; Additional File 1). Gene ontology analysis further confirms distinct enrichment in upregulated and downregulated biological processes in PoMiC compared to MiC (Figures. 1e and 1f; S1a). Genes associated with ribosomal biogenesis and translation show downregulation in PoMiC, indicative of anabolic repression, which includes purine metabolism and steroid biosynthesis. Simultaneously, genes involved in aerobic respiration and energy generation are upregulated in PoMiC, along with the *GLO*, *HSP*, *SNO*, and *SNZ* gene families, which are critical for survival under stress conditions (Additional File 1). Our results align with previous findings regarding the distinct biological signatures between MiC and PoMiC ^50,52,53^.

Similarly, PCA of transcriptomic profiles for samples treated with CR, RM, and combined CR+RM during both exponential and stationary phases distinctly separates them along PC1 and PC2 (Figure. 1c). Interestingly, CR and CR+RM samples cluster closely along PC1 and PC2, while RM-treated samples show a distinct deviation (Figure. 1c). These results suggest an intricate transcriptomic response in calorie-restricted and rapamycin-treated MiC and PoMiC. It is possible that CR and RM treatments may interact in a complex way. Some genes could be upregulated or downregulated in response to one treatment, and these genes may react differently when both treatments are used together. Additionally, other genes might have distinct responses when both treatments are combined. This interplay could contribute to the observed clustering. Furthermore, the differential separation of control and intervention groups in both exponential and stationary growth conditions highlights the distinct influence of CR, RM, and CR+RM treatments on the transcriptomic profiles in MiC and PoMiC. These observations are further supported by upset plots (Figures. S1b and S1c), which reveal that various conditions exhibit uniquely upregulated and downregulated differentially expressed genes (DEGs).

Next, our focus is to systematically analyze each intervention to gain a comprehensive understanding of why transcriptomic profiles of CR and CR+RM samples cluster closely, while RM-treated samples show deviation. This will enable us to uncover the specific genes and pathways involved in the distinct intervention conditions in MiC and PoMiC.

### Calorie restriction induced gene expression profiles in MiC and PoMiC

CR exerted a significant influence on gene expression, with its effects intricately linked to the growth phase. During the exponential phase, 62 genes exhibited significant upregulation, and 81 genes displayed significant downregulation in MiC (Figures. 2a; S2a; Additional File 2). These effects were more pronounced in the stationary phase, where 1583 genes were upregulated, and 1784 genes were downregulated in PoMiC (Figures. 2a; S2a; Additional File 2). Importantly, CR’s effects extended beyond the mere enhancement of transcriptomic profiles, as gene expression variations were evident across different growth phases. Specifically, 144 genes displayed unique differential expression in MiC, while 99 genes were shared between PoMiC (Figure. 2b; Additional File 2). Among these shared genes, 9 were consistently upregulated, and 41 were consistently downregulated (Figure. 2b), with 49 genes exhibiting divergent expression patterns (Figure. 2c).

**Figure 2.**
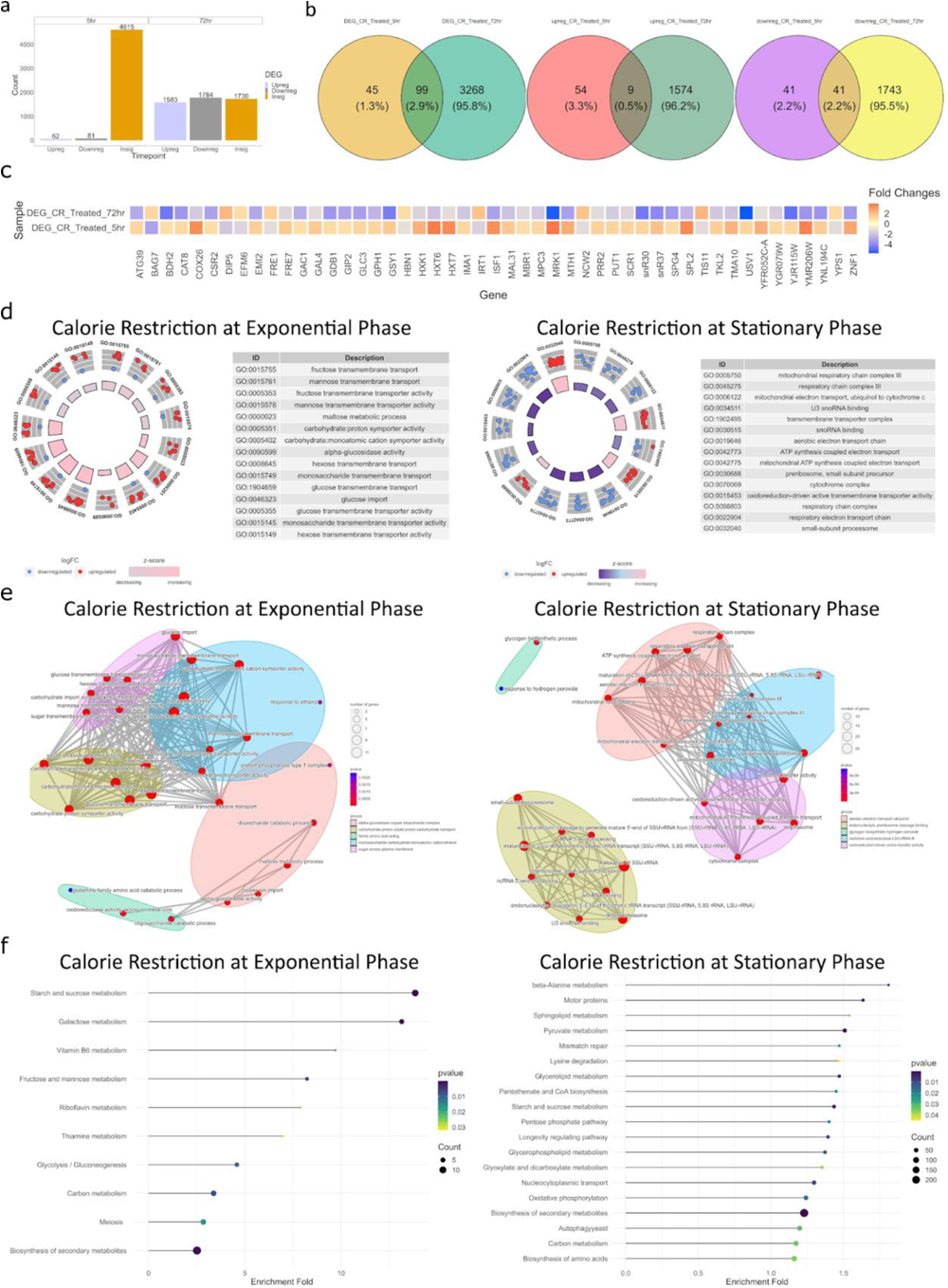
Calorie restriction-induced transcriptional changes in mitotic and postmitotic cells. (a) Bar plot illustrating the number of differentially expressed genes (DEGs) during both exponential (mitotic) and stationary (postmitotic) growth phases. Transcriptomic data from the calorie restriction treatment group is compared with vehicle control data for DEG analysis. Genes exhibiting an adjusted p-value ≤ 0.05 and an absolute log2 fold change ≥ 1 are considered as showing upregulated or downregulated in response to the treatment. (b) Venn diagrams showcasing the logical relationships between the results of DEGs analysis in the exponential and stationary phases. DEGs are further categorized into upregulated and downregulated groups, highlighting the shared and distinct gene expression responses to the treatments. The overlap and distinctness of DEGs in these growth phases are visually represented, highlighting shared and phase-specific gene expression changes. (c) Heatmap depicting genes that are regulated by the treatment interventions in both exponential and stationary phases yet exhibit differential expression patterns between the two growth phases. This visualization underscores the dynamic nature of gene expression responses to the treatments across different phases of growth. (d) Circular plots presenting the top gene ontology (GO) enrichments associated with DEGs in both exponential and stationary phases. The distinct biological processes and functions influenced by these gene expression changes are depicted, offering insights into the functional implications of the treatment interventions. (e) Network diagrams visually depict the enriched gene ontologies organized into distinct functional groups. These diagrams offer a holistic perspective on the interrelationships and interdependencies among gene functions that are impacted by the treatment interventions in both the exponential and stationary phases. (f) Dot plots showcasing the enriched KEGG pathways influenced by DEGs in the exponential and stationary phases. These plots offer a succinct overview of the key metabolic pathways and processes affected by the treatment interventions.

Biological processes involved in glucose import is upregulated including high-affinity glucose transporter *HXT* family genes such as *HXT2/4/6/7*, while the low-affinity glucose transporter *HXT1* exhibited downregulation (Figures. 2d and 2e; S2b and S2c). Concurrently, carbohydrate catabolism of alternative sugars is upregulated (Figures. 1d and 2e; S2b and S2c). Maltose metabolsim genes involved in high-affinity maltose transporters (alpha-glucoside transporter) *MAL11/31*, catabolism (alpha-D-glucosidase) *MAL12/32* and major isomaltase (alpha-1,6-glucosidase/alpha- methylglucosidase) *IMA1*, were significantly upregulated (Figures. 2d and 2e; S2b and S2c). Together, these analyses suggest that CR significantly influences carbohydrate metabolism and expresses compensatory biological processes in response to glucose deprivation. Through these processes, cells attained sufficient carbon sources for cellular processes including growth and metabolism (Figure. 2f).

At the stationary phase, biological processes involved in rRNA processing, including U3 small nucleolar RNA binding *UTP* family genes, were significantly upregulated (Figures. 2d and 2e; Figure. S2b). Conversely, mitochondrial respiratory-associated genes such as *COX13/5A/7*, *CYC7*, *CYT1*, and *QCR2/7/9/10* genes were downregulated (Figures. 2d and 2e; Figure. S2b). These findings suggest that PoMiC during CR treatment prioritize energy conservation and preventing the activation of mitochondrial function. Mitochondrial activation is linked to oxidative stress, which can affect the lifespan of cells ^54^.

Together, these processes contribute to diverse pathways, including glucose metabolism, amino acid biosynthesis, ribosome biogenesis, mismatch repair, autophagy, and longevity regulation (Figure. 2f; Figure. S2c). Overall, the transcriptomic signature of CR is evidently diverse between MiC and PoMiC growth conditions.

### Rapamycin induced gene expression profiles in MiC and PoMiC

While CR exhibited distinct effects on gene expression profiles in MiC and PoMiC cells, our investigation now focuses on the impacts of rapamycin treatment in these cell types. RM treatment consistently induced transcriptomic changes across both exponential and stationary growth phases. During the exponential phase, 31 genes were upregulated, and 200 genes were downregulated (Figures. 3a, S3a, and Additional File 3). This pattern was amplified in the stationary phase, with 1289 genes upregulated and 1590 genes downregulated (Figures. 3a, S3a, and Additional File 3). Among the 232 differentially expressed genes in MiC, 165 genes exhibited similar regulation in PoMiC, while 67 genes displayed unique regulation in MiC (Figure. 3b and Additional File 3). Notably, a bias towards downregulation was observed under RM treatment. Among genes consistently regulated in both MiC and PoMiC, 20 were persistently upregulated, 134 were consistently downregulated, and 11 displayed distinct expression patterns (Figure. 3c).

**Figure 3.**
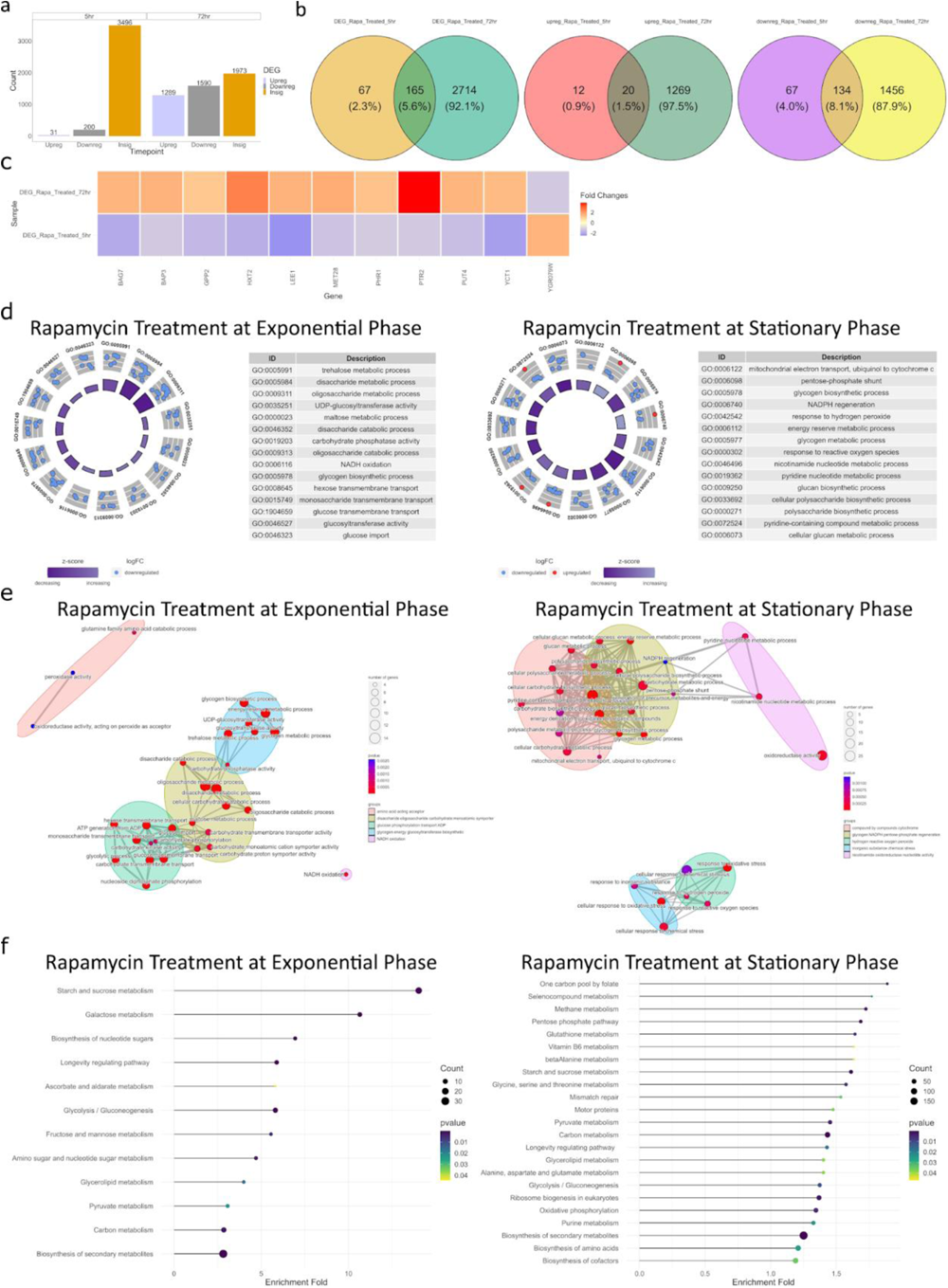
Rapamycin-induced transcriptional changes in mitotic and postmitotic cells. (a) Bar plot illustrating the number of differentially expressed genes (DEGs) during both exponential (mitotic) and stationary (postmitotic) growth phases. Transcriptomic data from the rapamycin treatment group is compared with vehicle control data for DEG analysis. (b) Venn diagrams showcasing the logical relationships between the results of DEGs analysis in the exponential and stationary phases. DEGs are further categorized into upregulated and downregulated groups, highlighting the shared and distinct gene expression responses to the treatments. The overlap and distinctness of DEGs in these growth phases are visually represented, highlighting shared and phase-specific gene expression changes. (c) Heatmap depicting genes that are regulated by the rapamycin treatment interventions in both exponential and stationary phases, yet exhibit differential expression patterns between the two growth phases. This visualization underscores the dynamic nature of gene expression responses to the treatments across different phases of growth. (d) Circular plots presenting the top GO enrichments associated with DEGs in both exponential and stationary phases. The distinct biological processes and functions influenced by these gene expression changes are depicted, offering insights into the functional implications of the rapamycin treatment interventions. (e) Network diagrams visually depict the enriched gene ontologies organized into distinct functional groups. These diagrams offer a holistic perspective on the interrelationships and interdependencies among gene functions that are impacted by the treatment interventions in both the exponential and stationary phases. (f) Dot plots showcasing the enriched KEGG pathways influenced by DEGs in the exponential and stationary phases. These plots offer a succinct overview of the key metabolic pathways and processes affected by the rapamycin treatment interventions.

Functional enrichment analysis of RM-treated MiC revealed a significant bias towards metabolism terms, particularly glucose metabolism during the exponential phase (Figures. 3d and 3e, S3b, and S3c). In yeast, glucose is the preferred carbon source, and its presence typically represses the utilization of alternative carbon sources. Surprisingly, maltose catabolic genes, including *MAL11/12/31/32* and *IMA1*, were downregulated in RM-treated MiC (Figure. S3b). This suggests that rapamycin may inhibit the utilization of maltose as a carbon source by the cells. Rapamycin’s downregulation of maltose metabolism genes could potentially shift the cellular preference even more towards glucose, inhibiting the yeast cells’ ability to efficiently utilize maltose.

Simultaneously, we found that genes involved in glycolytic intermediate pathways, including trehalose, pentose, mannose, and glycogen, were significantly downregulated alongside maltose catabolism (Figure. S3b). Glucose metabolism is interconnected with amino acid and lipid metabolism. Interestingly, amino acid and lipid biosynthesis processes are also downregulated in RM-treated MiC (Figure. S3c). The interplay of glucose metabolism with numerous cellular processes suggests that rapamycin treatment induces a broader metabolic reprogramming in MiC.

In the stationary phase, the pentose phosphate pathway, glycogen biosynthesis, redox processes, and glutathione metabolism were significantly affected in RM-treated PoMiC (Figures. 3d and 3e, S3b, and S3c). Genes involved in organonitrogen compound catabolic processes and mitochondrial functions were downregulated in RM-treated PoMiC (Figures. 3d and 3e, S3b, and S3c). Mitochondrial respiration generates reactive oxygen species (ROS), which are associated with the induction of cellular oxidative stress and DNA damage ^54^. Cells respond to various stresses by activating stress regulatory genes, including *CCP1* (cytochrome-c peroxidase), *CTT1* (Cytosolic catalase T), *DDR2* (DNA damage-responsive), and *DRE2* (response to DNA replication stress). Furthermore, genes involved in nicotinamide nucleotide metabolism and repair (*NNR1/2*, *NQM1*, and *PHO8*) were significantly downregulated (Figures. 3d and 3e, S3b, and S3c). Together, these findings suggest that rapamycin reduces metabolic activity and protects PoMiC against numerous stress conditions during the stationary growth phase to conserve energy and enhance damage repair over an extended timeframe.

### Comparative analysis of calorie restriction and rapamycin transcriptomic profiles in MiC and PoMiC

In our initial individual analysis of calorie restriction (CR) and rapamycin (RM) treatments, we gained insights into the transcriptomic changes induced by these interventions. Now, we seek to conduct a more thorough examination of the comparative transcriptomic profiles of CR and RM, shedding light on how they affect different cellular contexts, specifically in the MiC (exponential phase) and PoMiC (stationary phase).

A comparative analysis of DEGs between CR and RM interventions has revealed valuable insights into their shared and distinct impacts on gene regulation in MiC. During the mitotic growth phase, we observed that 52 genes were influenced by both treatments. Additionally, CR exclusively regulated 92 genes, and rapamycin uniquely controlled 179 genes (Figure. 4a; Additional File 4). Among these shared 52 genes, only two (*GFD2* and *YGR079W*) were upregulated, while 22 were downregulated in both treatments, underscoring the presence of 28 genes displaying divergent expression patterns (Figure. 4b). Notably, CR specifically upregulated genes associated with glucose metabolism, whereas rapamycin had the opposite effect, uniquely downregulating these genes (Figures. 4b and 4c; S4a and S4b; Additional File 5). While TORC1 is the target for both RM and CR ^55,56^, the pronounced divergence in transcriptomic profiles strongly indicates a clear distinction in their intervention effects on MiC.

**Figure 4.**
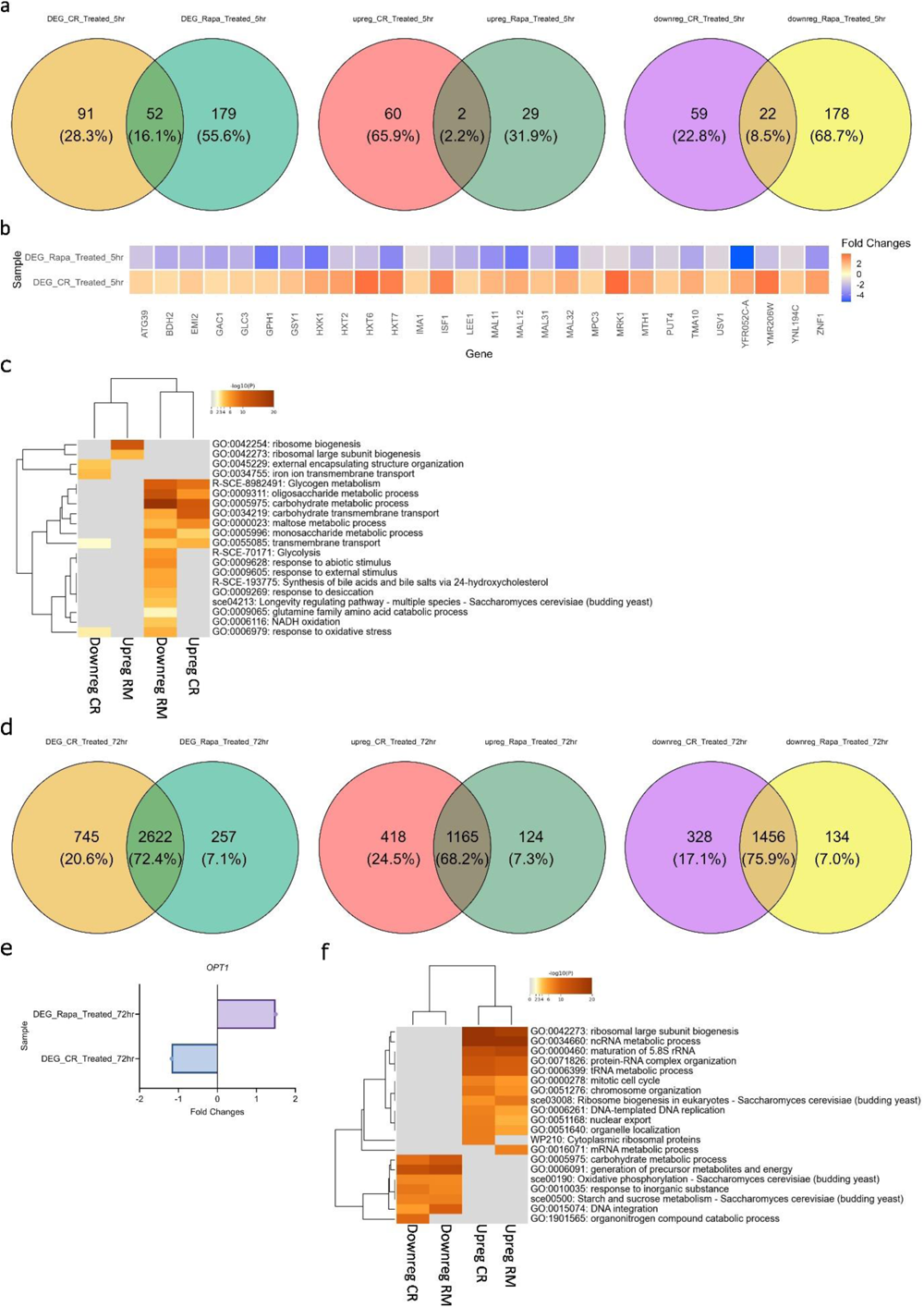
Comparative transcriptomic profiling of mitotic and postmitotic cells under calorie restriction and rapamycin treatments. (a and d) Venn diagrams depicting the comparison of differentially expressed genes (DEGs) between the calorie restriction and rapamycin treatments during (a) exponential (mitotic) and (d) stationary (postmitotic) growth phases. DEGs are further categorized into upregulated and downregulated groups, highlighting the shared and distinct gene expression responses to the treatments. (b and e) Heatmap showcasing DEGs that are commonly regulated by both calorie restriction and rapamycin treatments exhibit difference expression patterns during (b) exponential (mitotic), and (e) stationary (postmitotic) growth phases. (c and f) Functional enrichment analysis of upregulated and downregulated genes of both calorie restriction and rapamycin treatments during (c) exponential (mitotic), and (f) stationary (postmitotic) growth phases.

Moving to the stationary phase (PoMiC), we observed a greater similarity in transcriptomic profiles between CR and RM treatments, with 2622 genes shared (Figure. 4d; Additional File 4). Among these shared genes, 1165 were upregulated, and 1456 were downregulated, demonstrating some commonalities in their regulatory impact. Interestingly, only one gene, *OPT1*, exhibited discordant regulation, being downregulated by CR and upregulated by RM (Figure. 4e). These results suggest a potential convergence of effects between the treatments in PoMiC (Figures. 4f; S4c and S4d; Additional File 6), albeit with limited discordance in individual gene regulation.

Taken together, our comparative analysis underscores that RM is not a mere mimic of CR when considering differentially expressed genes (DEGs). While there is an overlap in DEGs between the two treatments, they also exert unique and distinct effects on gene expression in both MiC and PoMiC. This emphasizes the need for a nuanced understanding of the molecular mechanisms underlying CR and RM, particularly as they relate to different cellular contexts and phases of growth.

### Combined calorie restriction and rapamycin-induced gene expression profiles in MiC and PoMiC

The combined calorie restriction and rapamycin (CR+RM) treatment resulted in both shared and unique transcriptomic responses. During the exponential phase, 59 genes were upregulated, while 68 genes were downregulated in MiC (Figures. 5a and S5a, Additional File 7). However, in the stationary phase, 1633 genes were upregulated, and 1808 genes were downregulated in PoMiC (Figures. 5a and S5a, Additional File 7). Among the 128 genes differentially expressed in MiC, 91 genes were shared with PoMiC, while 37 genes displayed unique regulation in MiC (Figure. 5b). Within the shared 91 genes, 14 were consistently upregulated, and 35 were consistently downregulated (Figure. 5b), whereas 42 genes exhibited distinct expression patterns (Figure. 5c).

**Figure 5.**
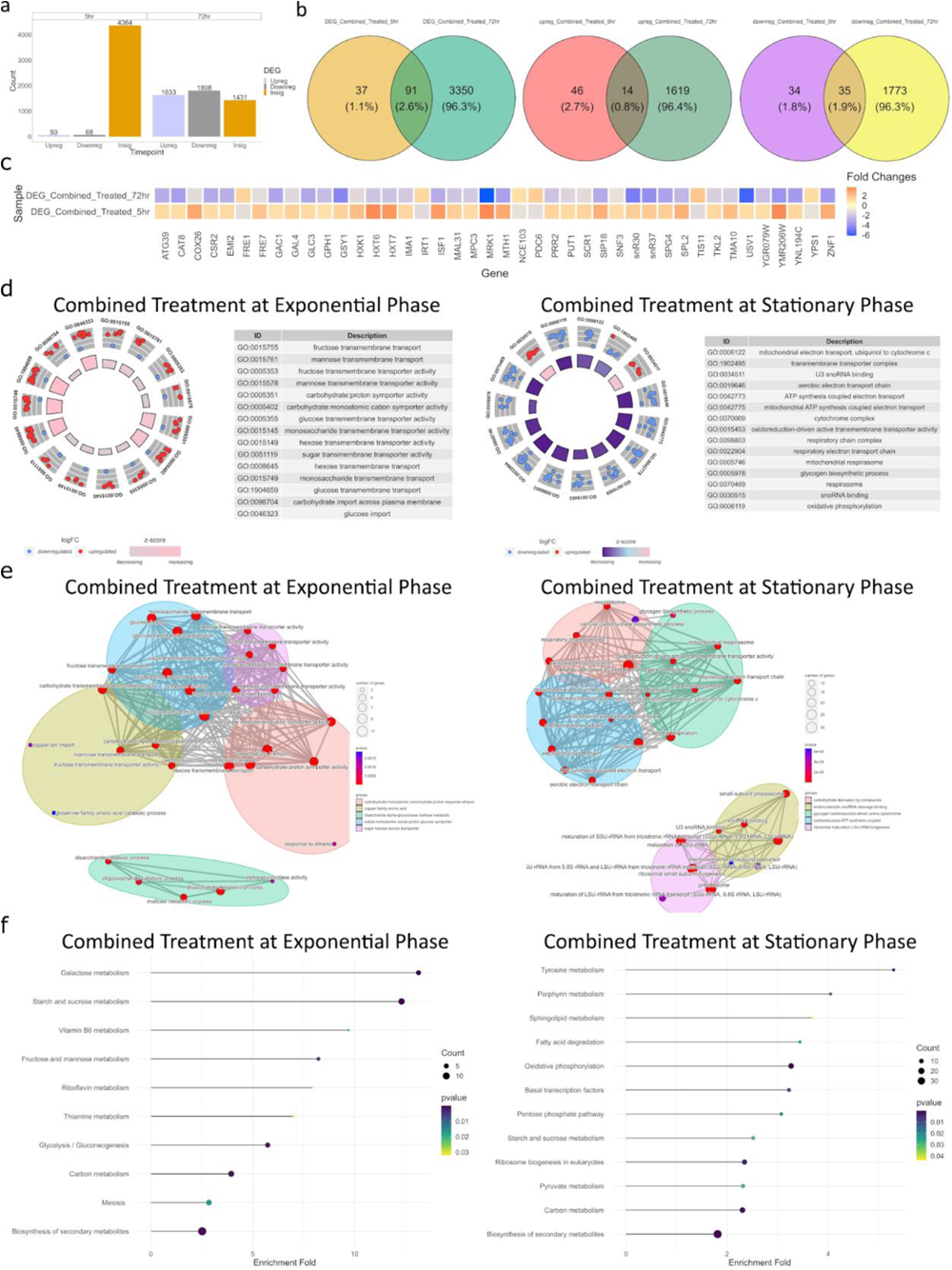
Combined calorie restriction and rapamycin-induced transcriptional changes in mitotic and postmitotic cells. (a) Bar plot illustrating the number of differentially expressed genes (DEGs) during both exponential (mitotic) and stationary (postmitotic) growth phases. Transcriptomic data from the combined treatment group is compared with vehicle control data for DEG analysis. (b) Venn diagrams showcasing the logical relationships between the results of DEGs analysis in the exponential and stationary phases. DEGs are further categorized into upregulated and downregulated groups, highlighting the shared and distinct gene expression responses to the treatments. The overlap and distinctness of DEGs in these growth phases are visually represented, highlighting shared and phase-specific gene expression changes. (c) Heatmap depicting genes that are regulated by the combined treatment interventions in both exponential and stationary phases, yet exhibit differential expression patterns between the two growth phases. This visualization underscores the dynamic nature of gene expression responses to the treatments across different phases of growth. (d) Circular plots presenting the top GO enrichments associated with DEGs in both exponential and stationary phases. The distinct biological processes and functions influenced by these gene expression changes are depicted, offering insights into the functional implications of the combined treatment interventions. (e) Network diagrams visually depict the enriched gene ontologies organized into distinct functional groups. These diagrams offer a holistic perspective on the interrelationships and interdependencies among gene functions that are impacted by the treatment interventions in both the exponential and stationary phases. (f) Dot plots showcasing the enriched KEGG pathways influenced by DEGs in the exponential and stationary phases. These plots offer a succinct overview of the key metabolic pathways and processes affected by the combined treatment interventions.

Furthermore, we conducted Gene Ontology (GO) analysis to uncover the biological processes modulated by CR+RM. Combinatorial interventions primarily influenced carbohydrate metabolism in MiC during the exponential phase (Figures. 5d-5f, S5b, and S5c). However, during the stationary phase, they affected ribosomal biogenesis and mitochondrial function in PoMiC (Figures. 5d-5f, S5b, and S5c). Interestingly, the functional profiles of combined CR+RM resembled those of CR alone (Figures. 5 and S5). These observations suggest that CR masks genetic expression and reveals the dominant epistasis interaction with RM in both MiC and PoMiC.

### Comparing combined calorie restriction and rapamycin-induced gene expression profiles with individual interventions in MiC and PoMiC

To gain insights into the genetic level, we compared the DEGs of CR, RM, and CR+RM. Remarkably, the impact of the combined treatment surpassed the cumulative effects of CR and RM. This was evident in the unique genes regulated by the combined treatment in MiC (Figures. 6a and 6b; Additional File 8) and PoMiC (Figures. 6c- and 6d; Additional File 8). During the exponential phase, the combined CR+RM treatment uniquely upregulated four genes (*VHR2*, *MLS1*, *SFC1*, *SIP18*) and downregulated four genes (*FIT3*, *PDC6*, *FRE5*, *MND1*) (Figure. 6a). Interestingly, an uncharacterized gene (*YGR079W*) was commonly upregulated in all three treatments (Figure. 6a). Another uncharacterized gene (*GFD2*), which was commonly upregulated between CR and RM treatments, was similarly upregulated in the combined CR+RM treatment (Figure. 6a), although the log-fold change was close to one (logFC: 0.944) (Additional File 7). These results suggest that the upregulation of the *YGR079W* and *GFD2* genes represents a common signature of CR, RM, and CR+RM interventions in MiC. Additionally, a single gene, *COS3*, was commonly upregulated by RM and CR+RM treatments, indicating that *COS3* regulation is predominantly controlled by the TORC1 pathway. We also found 47 overlapping genes that were differentially expressed in all treatments (Figure. 6b), and during the exponential phase in MiC, the CR+RM treatment exhibited greater similarity to CR than to RM (Figures. 6c; S6a and S6b; Additional File 9).

**Figure 6.**
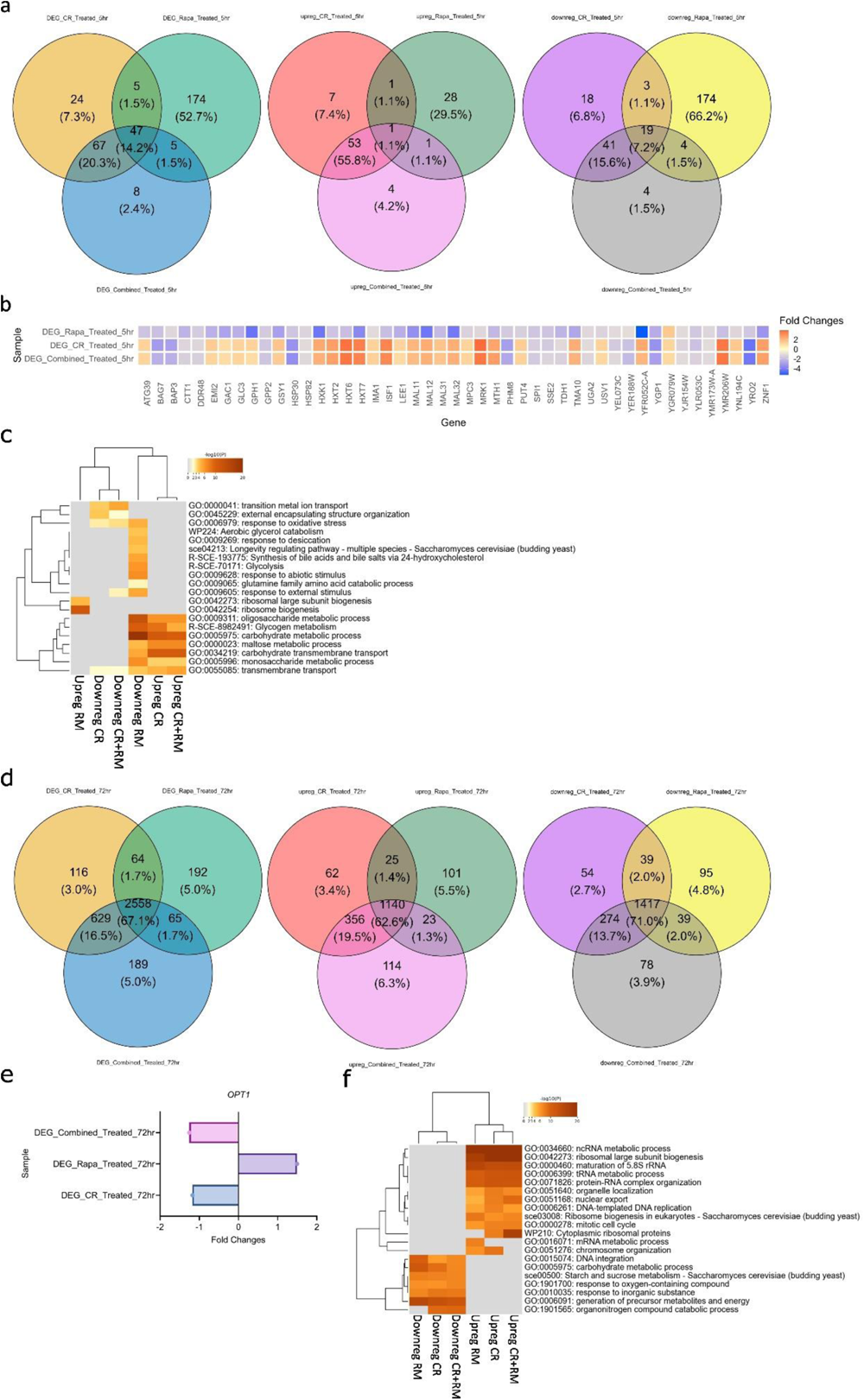
Comparative transcriptomic analysis of calorie restriction, rapamycin and combined treatments in mitotic and postmitotic cells. (a and d) Venn diagrams depicting the comparison of differentially expressed genes (DEGs) between the calorie restriction, rapamycin and combined treatments during (a) exponential (mitotic), and (d) stationary (postmitotic) growth phases. DEGs are further categorized into upregulated and downregulated groups, highlighting the shared and distinct gene expression responses to the treatments. (b and e) Heatmap showcasing DEGs that are commonly regulated by calorie restriction, rapamycin and combined treatments exhibit difference expression patterns during (b) exponential (mitotic), and (e) stationary (postmitotic) growth phases. (c and f) Functional enrichment analysis of upregulated and downregulated genes of calorie restriction, rapamycin and combined treatments during (c) exponential (mitotic), and (f) stationary (postmitotic) growth phases.

In the stationary phase, 2,558 genes exhibited common differential expression across all treatments. Notably, the *OPT1* gene, known for its upregulation in response to RM, displayed a significant downregulation upon exposure to the combined CR+RM treatment (Figure 6e). This observation highlights the presence of dominant epistatic regulation of the *OPT1* gene as a consequence of the CR intervention. Overall, the functional profiles of CR+RM treatment overlapped with those of CR and RM treatments in PoMiC (Figures 6f; S6c and S6d; Additional File 10). Nevertheless, the distinct regulation of genes by the combined treatment, separate from the effects of CR or RM individually (Figure 6d), suggests the possibility of a synergistic interaction between CR and RM interventions in PoMiC.

### Calorie restriction and rapamycin synergistically increase the lifespan of yeast and human postmitotic cells

CR and RM are well-known anti-aging interventions that can delay aging and extend the lifespan from yeast to human cells. Previous studies, including our own, have demonstrated that both interventions can extend the lifespan of yeast PoMiC ^11,57^. However, the combined effect of CR+RM has not been investigated before. Given that CR+RM treatment synergistically alters transcriptomics, we conducted experiments to assess their impact on cellular lifespan. We performed a series of postmitotic cellular lifespan (PoMiCL) assays using CR alone, RM alone, and a combination of both CR+RM during chronological aging in yeast. As expected, both CR and RM treatments independently increased lifespan (Figures. 7a-7c). Remarkably, the CR+RM combination treatment further extended the lifespan of PoMiC compared to individual treatments with either CR or RM (Figures. 7a-7c). These findings reveal that CR and RM synergistically increase the lifespan of PoMiC in yeast.

**Figure 7.**
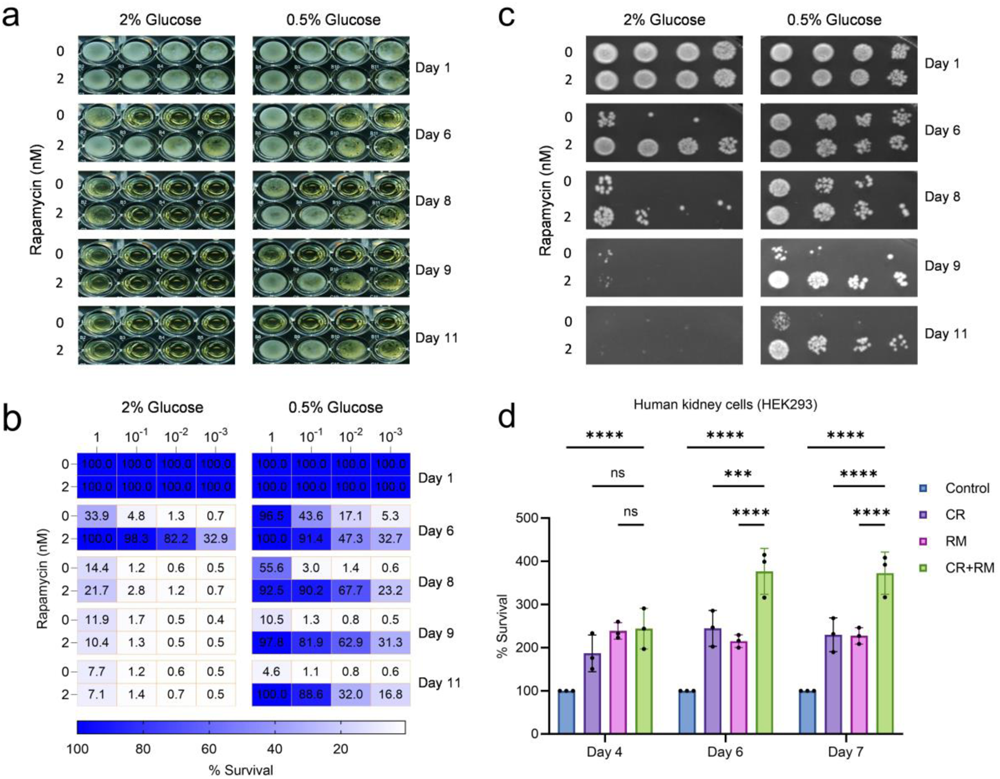
Enhancing longevity in yeast and human postmitotic cells through combined calorie restriction and rapamycin treatments. (a) The prototrophic yeast strain (CEN.PK113-7D) was grown in the synthetic defined medium under control (2% glucose), calorie restriction (0.5% glucose), 2 nM rapamycin and combined calorie restriction and rapamycin in 96-well plates at 30°C. The postmitotic cellular lifespan of aging culture was determined by the outgrowth survival method in YPD liquid medium. The stationary growth time point 72 hours was considered as Day 1. At various chronological time, aging culture was transferred to a second 96-well plate containing YPD medium. Outgrowth in 96-well plate was photographed after incubation for 24 hours at 30°C. (b) Outgrowth in YPD liquid medium was measured using a microplate reader. Survival was quantified relative to the outgrowth on day 1 and represented in a heatmap. (c) At various chronological time, 10-fold serial diluted aging cultures were spotted onto YPD agar plates. Outgrowth was photographed after incubation for 48 hours at 30°C. (d) Chronological lifespan of human kidney cells (HEK293) cells under postmitotic cellular condition was measured relative to the control outgrowth for the indicated day of analysis. Data are presented as means ± SD (n=3). Statistical significance was determined as follows: ***P < 0.001, and ****P < 0.0001, and ns (non-significant), based on a two-way ANOVA followed by Dunnett’s multiple comparisons test.

To broaden the scope of our findings, we investigated whether the synergistic interaction between CR and RM is conserved in human cells. We tested the effect of CR alone, RM alone, and the CR+RM combination in human kidney cells (HEK293) using the postmitotic chronological aging model. We found that the CR+RM combination significantly increased cellular lifespan more than CR or RM alone (Figure. 7d). In summary, these findings demonstrate that CR and RM interventions synergistically modulate transcriptomic profiles, leading to an increased lifespan in postmitotic cells, a phenomenon conserved across diverse species.

## DISCUSSION

In the past century, human life expectancy has witnessed a continuous upward trend, accompanied by a threefold surge in the global population from 2.5 billion in 1950 to nearly 7.8 billion in 2020, driven by advances in healthcare and evolving lifestyles ^58,59^. This demographic shift has resulted in a significant increase in the aging population, setting the stage for a rising prevalence of age-related diseases associated with dysfunction of mitotic and postmitotic cells.

In this study, we conducted a comprehensive analysis of transcriptomic signatures in yeast under mitotic and postmitotic growth conditions, with a particular focus on the effects of calorie restriction (CR), rapamycin (RM), and their combination (CR+RM) on gene expression profiles. Our findings shed light on the dynamic and intricate changes in gene regulation that occur in response to these interventions and provide valuable insights into their potential implications for cellular function and lifespan extension.

A striking revelation in our research is the clear segregation of transcriptomic profiles between mitotic cells (MiC) and postmitotic cells (PoMiC). This distinction is prominently showcased in the Principal Component Analysis (PCA), underscoring the substantial disparities in gene expression patterns between these two growth phases. This observation is consistent with prior studies ^50,52,53^ and underscores the significance of delineating the unique biological characteristics associated with different cellular growth conditions in yeast.

CR emerged as a potent modulator of gene expression, with its effects intricately linked to the growth phase of the cells. During the exponential growth phase, CR induced the upregulation of genes involved in glucose import and carbohydrate catabolism, suggestive of an adaptive response to glucose scarcity. In contrast, during the stationary phase, CR upregulated genes associated with rRNA processing while concurrently downregulating mitochondrial respiratory genes, indicating a shift toward energy conservation and stress mitigation. These findings illuminate the remarkable adaptability of yeast cells to nutrient limitations and highlight the fine-tuned regulatory mechanisms governing diverse growth phases.

RM treatment, on the other hand, exhibited a consistent and extensive impact on transcriptomic profiles across both exponential and stationary growth phases. Notably, RM exerted pronounced downregulation of genes related to glucose metabolism, glycolytic intermediates, amino acid biosynthesis, and lipid metabolism during exponential growth. This comprehensive metabolic reprogramming suggests a concerted effort to conserve energy resources. In postmitotic cells, RM-influenced pathways such as the pentose phosphate pathway, glycogen biosynthesis, and redox processes, hint at a strategy aimed at enhancing stress resistance and facilitating damage repair. These findings underscore the multifaceted nature of RM’s influence on cellular metabolism and stress response.

One of the most notable findings in our study is the unequivocal demonstration that RM is not a mere mimic of CR. While both interventions share some commonalities in their effects on gene regulation, they also manifest distinctive and independent impacts on the transcriptomic landscape. This divergence in gene expression patterns serves as a resounding reminder that RM operates through its own intricate pathways, apart from those influenced by CR. This finding has profound implications for our understanding of how these interventions impact cellular physiology. The difference in gene expression responses between CR and RM interventions was previously reported in the liver and skeletal muscle ^55,56^. Our data support these findings and further elucidate the largely distinct mechanisms by which these interventions act in a cellular context. Together, these findings show that these differences are conserved throughout evolutionary species from yeast to humans. The combination of CR+RM resulted in unique transcriptomic profiles that surpassed the cumulative effects of CR and RM alone. These findings deepen our understanding of the complex regulatory mechanisms particularly in PoMiC, indicating a potential epistatic interaction between the two interventions.

Highlighting some of the noteworthy genes from our study, including *OPT1*, *YGR079W*, *GFD2*, and *COS3*, is indeed worthwhile. *OPT1* is a gene with multiple aliases, including *HGT1*, *GSH11*, and *DUG4*, highlighting its versatile roles in transporting molecules like glutathione and phytochelatin, showcases contrasting regulation by CR and RM, hinting at a complex interplay between nutrient sensing and stress response pathways. *YGR079W*, an uncharacterized gene consistently upregulated by CR, RM, and CR+RM, presents an exciting frontier for research, potentially unveiling novel mechanisms of cellular adaptation and stress response. *GFD2*’s shared upregulation by CR and RM signifies a common signature of gene regulation in response to nutrient restriction and TORC1 inhibition, offering insights into pathways promoting healthy aging. Lastly, *COS3*, predominantly regulated by the TORC1 pathway, emphasizes TORC1’s pivotal role in protein turnover and cellular homeostasis. These genes collectively hold promise for future aging research, where unraveling their functions and interactions may uncover crucial pathways influencing stress adaptation, longevity, and age-related diseases, offering exciting prospects for novel therapeutic strategies.

The most intriguing finding of our study is the synergistic interaction between CR and RM, particularly in extending the lifespan of postmitotic cells. This phenomenon was observed in both yeast and human cells, highlighting its potential conservation across species. The combination of CR+RM resulted in unique transcriptomic profiles that surpassed the cumulative effects of CR and RM alone. These profiles revealed a convergence of effects, particularly in PoMiC, indicating a potential epistatic interaction between the two interventions.

Additionally, to substantiate the intriguing findings regarding the unique gene regulation patterns induced by the combined calorie restriction and rapamycin (CR+RM) intervention, we conducted further validation through lifespan experiments. Our study provided compelling evidence that CR and RM individually extended the lifespan of yeast postmitotic cells. However, the pivotal question remained: Does the combined CR+RM intervention exhibit a synergistic effect on lifespan extension? The answer was affirmative. The CR+RM combination treatment was found to synergistically increase the lifespan compared to either CR or RM alone. This validation not only reinforces the distinct impact of CR+RM on gene regulation but also underscores its potential translational relevance. Further, the cross-species validation in human cells highlights the potential for CR+RM to influence longevity across diverse organisms and provides a promising avenue for future research on its application in human health.

In conclusion, the unique gene regulation observed with the CR+RM intervention presents an exciting opportunity for further investigation. Analyzing the functions, interactions, and implications of these genes may advance our understanding of aging processes and offer potential strategies to promote unraveling the complexity of cellular aging. It is worth noting that our study not only sheds light on the distinctive effects of CR+RM but also provides initial insights into the independent impacts of CR and RM. Alone, CR and RM each exhibited distinct regulatory effects on gene expression patterns, emphasizing the importance of careful consideration regarding the choice of intervention in different cell types and physiological conditions. CR appears to influence gene expression by promoting metabolic adaptation and stress resistance, whereas RM exhibits regulatory effects linked to nutrient sensing and cellular growth inhibition. The combination of these interventions, as demonstrated in our study, unveils a unique interplay of these pathways, offering a potential paradigm shift in our approach to understanding and modulating aging and longevity across diverse species. This research sets the stage for further exploration into the molecular mechanisms governing the effects of CR, RM, and their synergistic combination, ultimately paving the way for innovative strategies to enhance healthspan and delay the onset of age-related diseases.

## METHODS

### Yeast strain cultivation

The prototrophic *S. cerevisiae* CEN.PK 113-7D Matα strain ^51^ was revived from a glycerol stock and cultured on YPD agar medium (1% Bacto yeast extract, 2% Bacto peptone, 2% glucose, and 2.5% Bacto agar) for 2-3 days at 30°C. A single yeast colony was isolated and cultivated in a conical flask containing synthetic defined (SD) medium (6.7 g/L yeast nitrogen base with ammonium sulfate without amino acids and 2% glucose) overnight at 30°C with shaking at 220 rpm. Subsequently, an overnight culture was inoculated into fresh SD medium with an initial optical density of 0.2 (OD600nm). The culture flasks were then incubated at 30⁰C with continuous shaking at 220 rpm throughout the duration of the experiments.

### Cell culture chemical exposure

Stock solution of rapamycin was prepared in dimethyl sulfoxide (DMSO). The final concentration of DMSO did not exceed 1% in yeast and 0.01% in human cell lines experiments.

### RNA extraction

RNA extraction was performed following established protocols ^60^. Total RNA was extracted from yeast cells using the Qiagen RNeasy mini kit. Yeast cultures were sampled at various time points, and the initial step involved mechanically disrupting the cells according to the manufacturer’s guidelines. Subsequently, the concentration and integrity of the RNA were assessed using the ND-1000 UV-visible light spectrophotometer from Nanodrop Technologies and the Bioanalyzer 2100 with the RNA 6000 Nano Lab Chip kit by Agilent.

### RNA sequencing

High-quality RNA samples were subsequently prepared for paired-end RNA sequencing by the Novogene facility. An enrichment process targeting the polyadenylated mRNA fraction was employed. Subsequent to the enrichment step, complementary DNA (cDNA) libraries were generated from the enriched mRNA through reverse transcription. Bioanalyzer analysis was employed to validate the quality and size distribution of the cDNA libraries. The actual RNA sequencing process was performed using the NovaSeq PE150 platform.

### Data processing and alignment

The raw sequencing reads underwent alignment using the nf-core RNAseq pipeline (nf-core/rna-seq revision 3.8.1), which employed the STAR aligner and RSEM quantification ^61^. The alignment process utilized the *S. cerevisiae* reference genome with a matching annotation (GTF format ver. 1.105) sourced from Ensembl, resulting in the generation of gene-level expression counts ^62^.

### Differential gene expression analysis

To assess the impact of the CR, RAPA, and Combined treatments on gene expression profiles, an in-depth differential gene expression (DEG) analysis was conducted. Gene expression counts were subjected to normalization, and differential expression was evaluated through the utilization of DESeq2 ^63^. This analysis encompassed intra-group comparisons both before and after the diauxic shift, as well as inter-group comparisons across the distinct treatment groups.

### Functional annotation and enrichment analysis

Significantly differentially expressed genes underwent functional annotation and enrichment analyses. These analyses identified gene ontology (GO) terms, biological pathways, and molecular functions, and were carried out using the clusterProfiler package within the R environment ^64,65^ and metascape analysis online tool ^66^. This step provided valuable insights into the biological processes influenced by the interventions.

### Analysis of postmitotic cellular lifespan of yeast during chronological aging

The lifespan of yeast postmitotic cells was evaluated through a chronological aging study, following a previously established methods ^57^. Yeast cultures were initially grown in synthetic defined (SD) medium at 30°C with 220 rpm agitation overnight. Subsequently, these cultures were diluted to an initial optical density at 600nm (OD600nm) of 0.2 in fresh SD medium. This was done under three distinct experimental conditions: calorie restriction (CR), Rapamycin (RM) treatment, and a combination of both (CR+RM). These conditions were used to initiate the postmitotic cellular lifespan (PoMiCL) experiment. In brief, the yeast cell cultures were cultivated in 96-well plates and allowed to enter the postmitotic phase at 30°C. At various chronological time points, the survival of aging postmitotic cells was assessed using two methods: (i) outgrowth in YPD liquid medium and (ii) outgrowth on YPD agar medium through a spotting assay. For the YPD liquid medium outgrowth method, yeast stationary cultures (2 μL) at different ages were transferred to a second 96-well plate containing 200 μL of YPD medium. This plate was then incubated for 24 hours at 30°C, and the outgrowth was quantified by measuring the OD600 using a microplate reader. Similarly, for the YPD agar medium outgrowth method, 10-fold serial diluted aging cultures spotted (3 μL) onto YPD agar plates and incubated for 48 hours at 30°C. The outgrowth on the YPD agar plate was documented through photography using the GelDoc imaging system.

### Analysis of postmitotic cellular lifespan analysis of human cells during chronological aging

To determine the longevity of postmitotic human cells as they age chronologically, we employed a propidium iodide (PI) fluorescent-based method to measure cell death. In this study, 80,000 cells were seeded in a 96-well culture plate with a D10 medium, maintaining a non-dividing state. The cells were subjected to various conditions, including calorie restriction (CR), Rapamycin (RM) treatment, and a combination of both (CR+RM). The incubation was carried out at 37°C in an environment with 5% CO2. Throughout different time points representing various stages of chronological aging, we monitored cell survival by quantifying PI fluorescence. This was achieved using a microplate reader, which excited the PI at 535 nm and measured the emitted fluorescence at 617 nm.

### Statistical analyses and figure preparation

Statistical analyses were performed using the R software (version 4.31), pertinent bioinformatics packages and GraphPad Prism v.10 software. The reported p-values are two-tailed, with statistical significance determined using predetermined thresholds (e.g., adjusted p < 0.05). Figures were generated within the R environment, utilizing packages including ggplot2, UpSetR, ggvenn, GOplot ^67–69^, metascape analysis online tool ^66^ and GraphPad Prism v.10 software. These tools facilitated the creation of informative visualizations as well as graphical representations of the results obtained.

## ACKNOWLEDGMENTS

This work was supported by Bioinformatics Institute (BII), A*STAR Career Development Fund (C210112008) and the Healthy Longevity Catalyst Awards grant (MOH-000758–00).

## AUTHOR CONTRIBUTIONS STATEMENT

YZ, AN, TINC, JLJ, and MA conducted the experiments. YZ and MA wrote the manuscript. MA conceived and supervised the project. All authors have read the manuscript.

## SUPPLEMENTARY FIGUREURE LEGENDS

**Figure S1.**
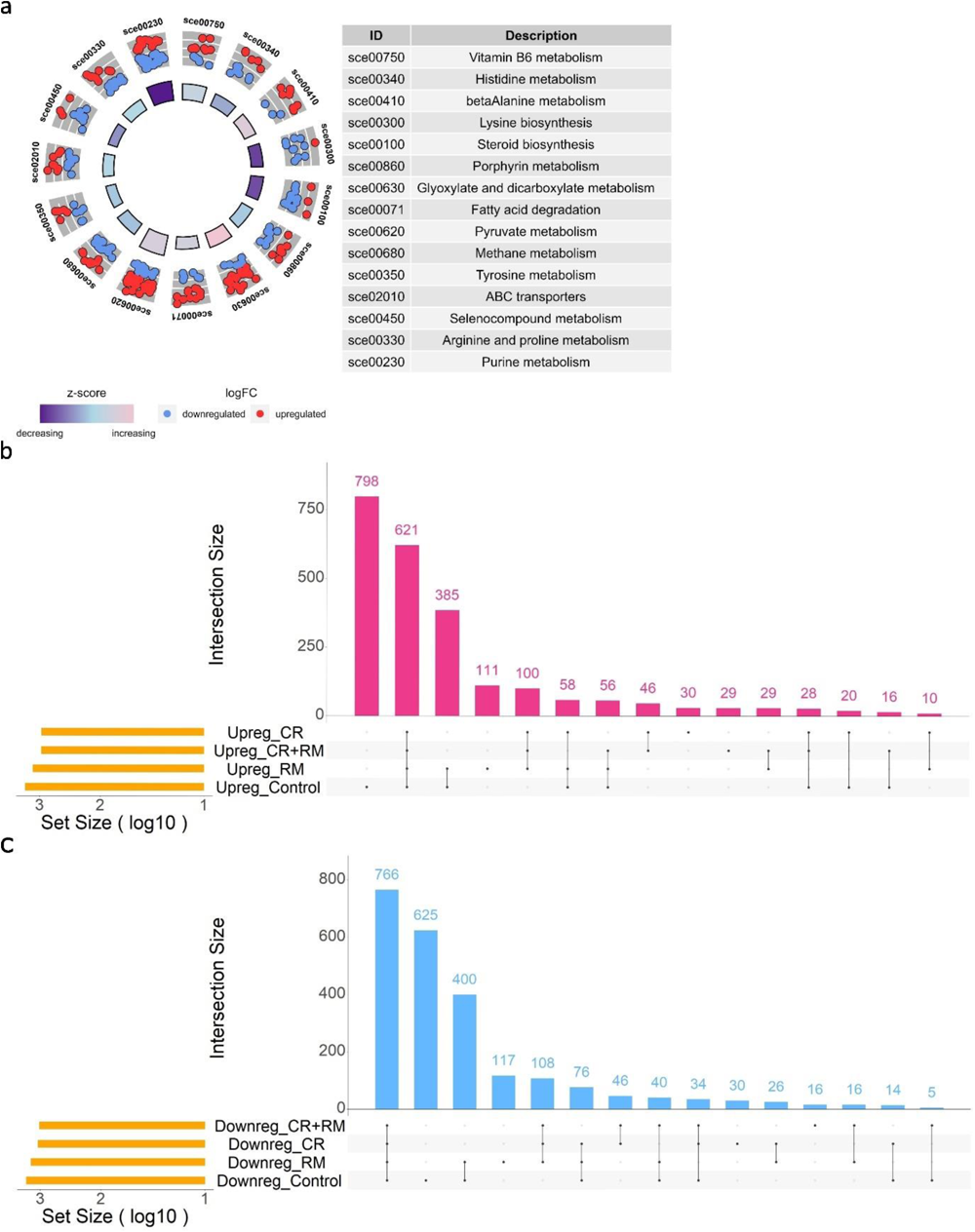
Transcriptomic analysis of exponential phase culture and stationary phase culture across all treatments, Related to Figure 1. (a) Circular plots presenting the top KEGG enrichments associated with DEGs in control sample for stationary phase postmitotic cells compared to exponential phase mitotic cells. (b and d) Upset plot illustrating the comparison of (a) upregulated genes, and (b) downregulated genes from control, calorie restriction (CR), rapamycin (RM) and combined (CR+RM) treatments for stationary phase postmitotic cells compared to exponential phase mitotic cells.

**Figure S2.**
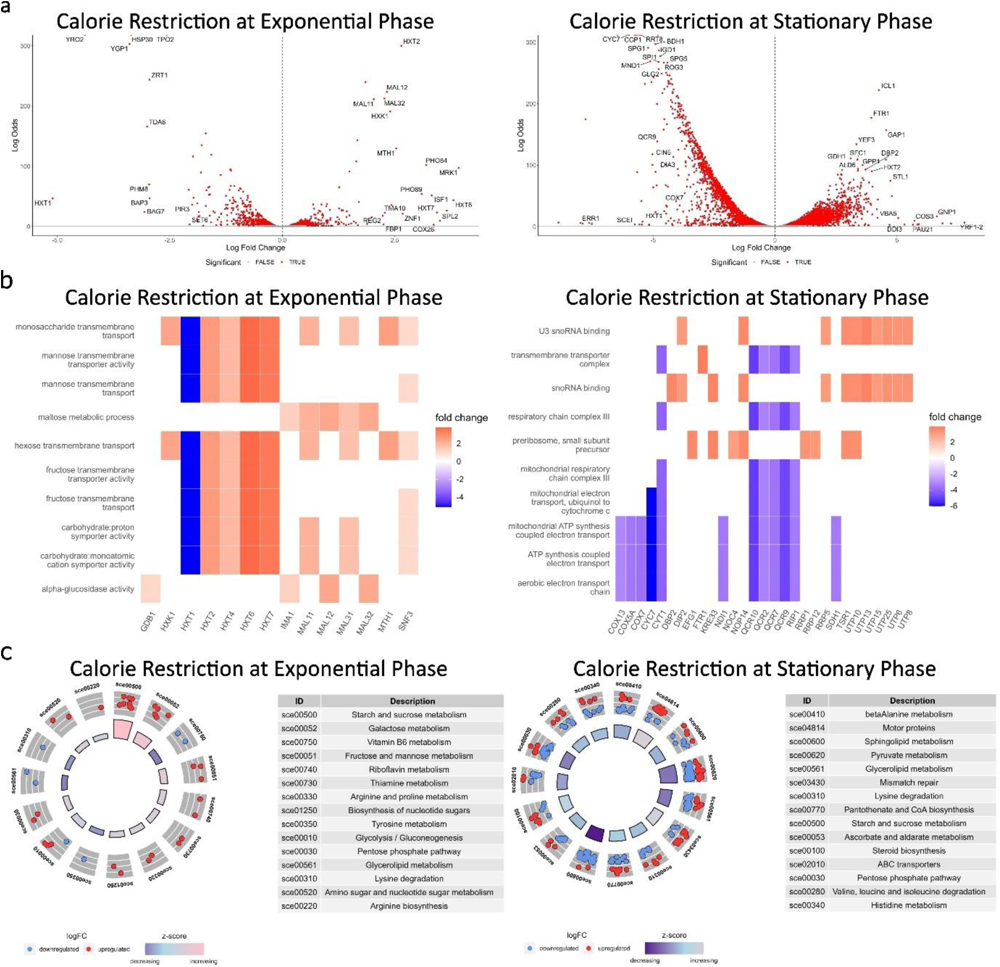
Transcriptomic analysis of calorie restriction treatment in mitotic and postmitotic cells, Related to Figure 2. (a) Volcano plots illustrating the distribution of differentially expressed genes (DEGs) resulting from calorie restriction treatment during both exponential (mitotic) and stationary (postmitotic) growth phases. (b) Heatmap illustrating enriched gene ontologies and the genes responsible for regulating multiple processes. These diagrams provide a comprehensive view of the interconnectedness of gene functions influenced by the treatment interventions during both exponential and stationary phases. (c) Circular plots presenting the top KEGG enrichments associated with DEGs in both exponential and stationary phases.

**Figure S3.**
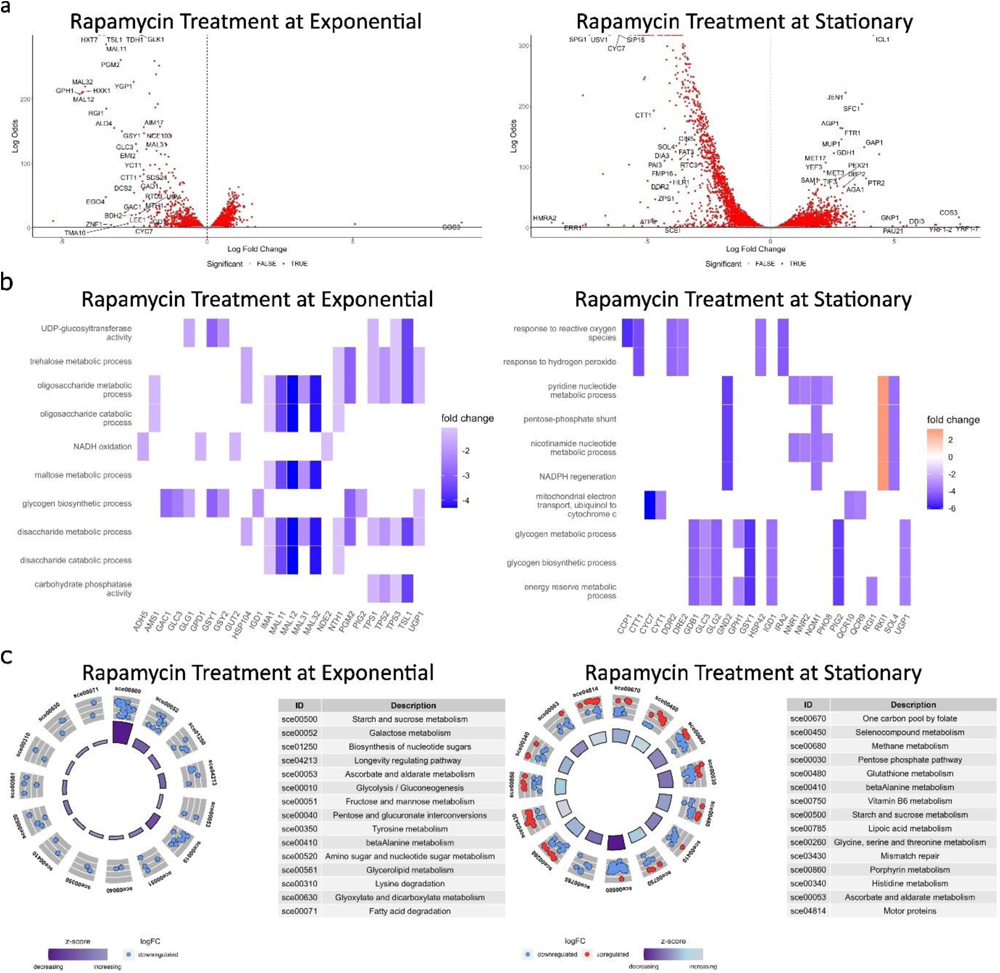
Transcriptomic analysis of rapamycin treatment in mitotic and postmitotic cells, Related to Figure 3. (a) Volcano plots illustrating the distribution of differentially expressed genes (DEGs) resulting from rapamycin treatment during both exponential (mitotic) and stationary (postmitotic) growth phases. (b) Heatmap illustrating enriched gene ontologies and the genes responsible for regulating multiple processes. These diagrams provide a comprehensive view of the interconnectedness of gene functions influenced by the treatment interventions during both exponential and stationary phases. (c) Circular plots presenting the top KEGG enrichments associated with DEGs in both exponential and stationary phases.

**Figure S4.**
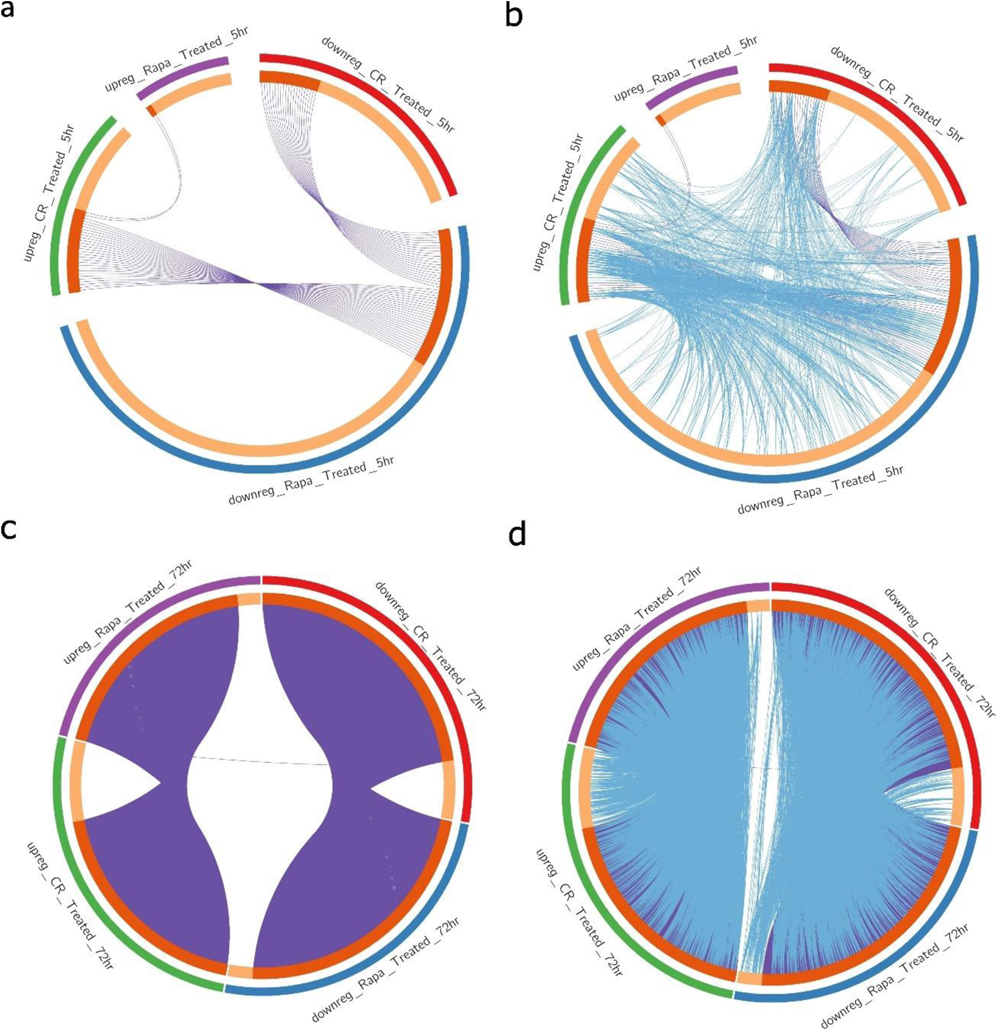
Comparing the transcriptomic profiles of calorie restriction and rapamycin in mitotic and postmitotic cells, Related to Figure 4. (a and c) The metascape circos plots show the overlapping genes from the input differentially expressed genes (DEGs) of both calorie restriction and rapamycin treatments during (a) exponential (mitotic), and (c) stationary (postmitotic) growth phases. See also Additional File 5. (b and d) The metascape circos plots show the overlapping genes and functional enrichment from the input differentially expressed genes (DEGs) of both calorie restriction and rapamycin treatments during (a) exponential (mitotic), and (c) stationary (postmitotic) growth phases. See also Additional File 6.

**Figure S5.**
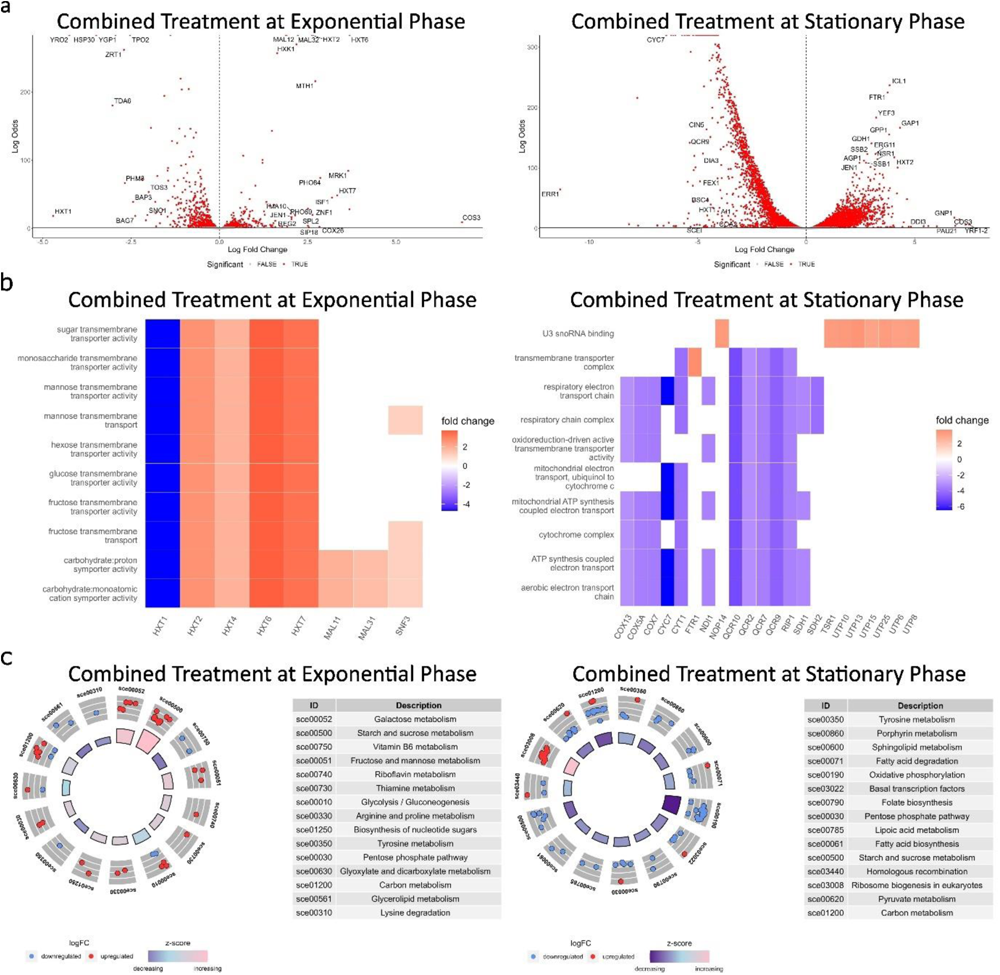
Transcriptomic analysis of combined treatment in mitotic and postmitotic cells, Related to Figure 5. (a) Volcano plots illustrating the distribution of differentially expressed genes (DEGs) resulting from combined treatment during both exponential (mitotic) and stationary (postmitotic) growth phases. (b) Heatmap illustrating enriched gene ontologies and the genes responsible for regulating multiple processes. These diagrams provide a comprehensive view of the interconnectedness of gene functions influenced by the treatment interventions during both exponential and stationary phases. (c) Circular plots presenting the top KEGG enrichments associated with DEGs in both exponential and stationary phases.

**Figure S6.**
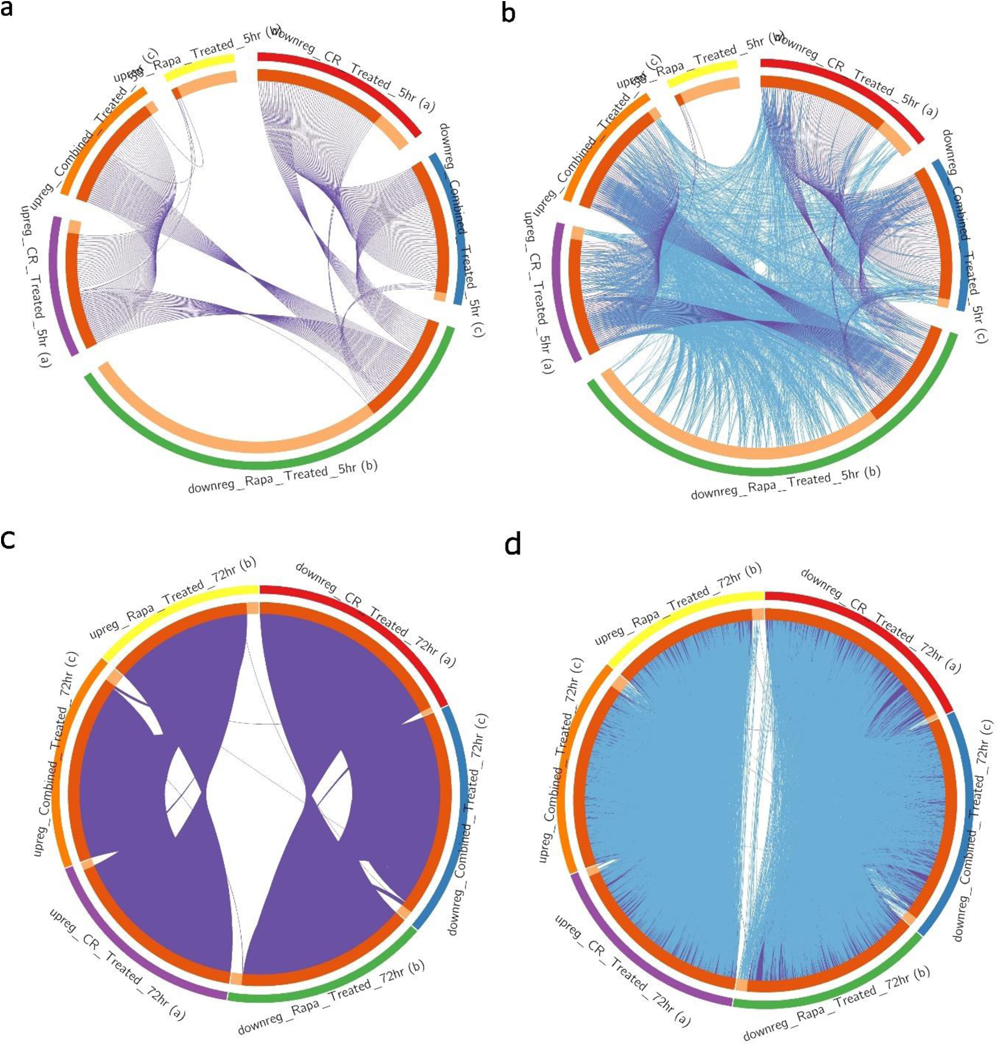
Comparing the transcriptomic profiles of calorie restriction, rapamycin and combined treatments in mitotic and postmitotic cells, Related to Figure 6. (a and c) The metascape circos plots show the overlapping genes from the input differentially expressed genes (DEGs) of calorie restriction, rapamycin and combined treatments during (a) exponential (mitotic), and (c) stationary (postmitotic) growth phases. See also Additional File 9. (b and d) The metascape circos plots show the overlapping genes and functional enrichment from the input differentially expressed genes (DEGs) of calorie restriction, rapamycin and combined treatments during (a) exponential (mitotic), and (c) stationary (postmitotic) growth phases. See also Additional File 10.

